# Genetic connectivity and diversity of endangered species the scalloped hammerhead shark *Sphyrna lewini* (Griffith & Smith 1834) population in Indonesia and Western Indian Ocean

**DOI:** 10.1101/2020.03.10.985465

**Authors:** Sutanto Hadi, Noviar Andayani, Effin Muttaqin, Benaya M Simeon, Muhammad Ichsan, Beginer Subhan, Hawis Madduppa

## Abstract

The scalloped hammerhead shark *Sphyrna lewini* is an endangered species which expected to population declined worldwide including in Indonesia due to overexploited. However, there is a lack of information regarding recent population structure to promote proper management and conservation status in Indonesia. This study aimed to investigate the genetic diversity, population structure and connectivity of *S. lewini* population in Indonesia from three major sharks landing sites in Aceh (n= 41), Balikpapan (n= 30), Lombok (n= 29), and additional sequences retrieved from West Papua (n= 14) and Western Indian Ocean population (n= 65). Analyses of mitochondrial CO1 gene successfully identified a total of 179 sequences of *S. lewini* with an average 594 bp nucleotide with 40 polymorphic loci in 4 haplotypes for Indonesian population and 8 haplotypes for Western Indian Ocean. The overall values of genetic diversity in Indonesia was high (Hd= 0.7171; π= 0.0126), with the highest was in Aceh (Hd= 0.6683; π= 0.0198), and the lowest was in Papua (Hd= 0.1429; π= 0.0005), while in Western Indian Ocean the overall value was fairly low (Hd= 0.2322; π= 0.0010). The AMOVA and *F*_ST_ revealed three significant population subdivisions in Indonesia (*F*_ST_= 0.4415; *p* < 0.001) with separated population for Aceh and West Papua, and a mixing population between Balikpapan and Lombok (*F*_ST_= 0.044; *p* = 0.089), whereas relatively no significant differentiation within population in Western Indian Ocean (*F*_ST_= −0.0131; *p* = 0.6011), and significant different level showed by Indonesian population compared with Western Indian Ocean population (*F*_ST_= 0.7403; *p* < 0.001). The construction of haplotype network exhibited evidence of gene flow and haplotype sharing between populations. This result indicated a complex and limited connectivity population of *S. lewini* in Indonesia, and between Western Indian Ocean in regional scale which need co-management action across region.

## Introduction

The scalloped hammerhead shark, *Sphyrna lewini* (*S. lewini*) is semi-oceanic species which distributed circumglobally along tropical waters, abundant both around continental margins and close to the coastal region and islands [1–2]. This species has a unique modification of laterally expanded head which enhanced them with the ability to navigate and follow geomagnetic orientation across the ocean [3–5]. *S. lewini* has a capacity to move in high rates of dispersal. Female *S. lewini* exhibits site fidelity to single nursery area and indicate no evidence of continuing inter oceanic migration, otherwise male *S. lewini* spread in a long distance spatial across ocean waters with clear evidence in cross reproduction and gametes transmission [6].

*S. lewini* is one of the threatened species of sharks which estimated one to three million were killed annually worldwide due to fishing and shark fins trade. This species was considered to be an underexploited species in 1999, however, currently listed to be endangered species in the IUCN red list [7] and Appendix II of The Convention on International Trade in Endangered Species (CITES). High exploitation of the population of *S. lewini* may has an impact on the population structure, reducing the fecundity of species and genetic diversity of populations [8]. Furthermore, this species has low resilience to exploitation since late sexual maturity and a long lifetime in nature [9–10]. The study of population genetic have become an important tool for understanding population connectivity, defining fisheries management and conservation strategies [11]. Genetic of population structure of *S. lewini* which important to fisheries stock management has been widely investigated in different coastal area and ocean basin on a global and regional scale [6, 12]. Duncan *et al*. [13] revealed a global phylogeographic study of *S. lewini* which indicated that Indo-West Pacific was a centre of diversity for tropical sharks like *S. lewini* with a high-unique genetic diversity, however, no samples from Indonesia were included, also the study by Ovenden *et al.* [14] which conducted with limited samples from Indonesia.

The aim of this study was to investigate genetic diversity, population structure, and connectivity of *S. lewini* among population in Indonesia where the population are affected by fishing activities, observe connectivity in regional scale in Western Indian Ocean, and also suggest the implication for the species management and conservation.

## Material and Methods

### Ethics Statement

All specimens were collected in dead condition, no live specimens were obtained. Therefore, no approval from any institutional animal ethics committee was required. The samples collection and transport were in accordance with the regulation of Ministry of Marine Affairs and Fisheries of the Republic of Indonesia Number 5/PERMEN-KP/2018 on the prohibition of cowboy shark and hammerhead shark export from Indonesia. This study was also approved under permission number 276/BPSPL.03/PRL/X/2018 and 319/PNK/BPSPL.03/PK.230/REKOM/X/2018 by Ministry of Marine Affairs and Fisheries of the Republic of Indonesia.

### Tissue samples collection

A total of 100 tissue samples of *S. lewini* were collected from October 2017 to November 2018, consisting of 41 samples from Meulaboh and Aceh Jaya fishing port (Aceh), 30 samples from local shark landing in Balikpapan, and 29 samples from Tanjung Luar fishing port (Lombok). The samples were dissected from flesh or fin tissues approximately 0.5 cm^3^ preserved in sample bottle contained 96% ethanol.

### DNA extraction, amplification and sequencing

Extraction of DNA conducted in Biodiversity and Biosystematics Laboratory, IPB University following the protocol of the gSYNC DNA extraction kit product. A fragment of mitochondrial Cytochrome Oxidase subunit-I gene (COI) was amplified using forward primer fish-BCL 5’ TCA ACY AAT CAY AAA GAT ATY GGC AC ‘3 and reverse primer fish-BCH 5’ACT TCY GGG TGR CCR AAR AAT CA ‘3 [15–16] (Baldwin *et al*. 2009; Madduppa *et al.* 2016) in a 24 μL reaction mixture consisted of 3 μL DNA template, 12.5 μL MyTaq HS Red Mix, 9 μL ddH_2_O, 1.25 μL forward and reverse primer respectively. Mixture product was processed in a polymerase chain reaction (PCR) thermocycler following a modified cycle [17–18] (Prehadi *et al.* 2015; Hadi *et al.* 2019). The step was initiated with pre denaturation at 94 °C for 15 s, followed by 38 cycles of denaturation at 94 °C for 30 sec, annealing at 50 °C for 30 sec, extension at 72 °C for 45 sec, and finished with a final extension at 72 °C for 10 min. The amplicons were visualized by gel electrophoresis agarose 1.5 % smeared with ethidium bromide at 100 V for 20 min. The gel was observed under UV light to determine a band indicating the presence of DNA fragments. Sequence reaction was performed using the BigDye Terminator v3.1 cycle sequencing kit with an optimized protocol of Applied Biosystems 3730 genetic analyser at the 1st BASE, Malaysia.

### Data Analysis

#### Genetic diversity

Sequence of 100 mitochondrial DNA CO1 were edited and aligned using the CrustalW algorithm implemented in MEGA 6.06 [19]. Genetic diversity parameters including the number of haplotypes, haplotype diversity (Hd), and nucleotide diversity (π) were calculated using the DNASp v6 [20] and Arlequin v.3.5 program [21]. We also reanalysed additional COI sequences data of *S. lewini* retrieved from GenBank (Table 1) from West Papua (n= 14) obtained by Sembiring *et al.* [22] and sequences from previous study in India (n= 6) [23], United Arab Emirates (n= 30) [24], and Madagascar (n= 29) [25] to show genetic diversity in Indonesian and Western Indian Ocean.

**Table 1.**
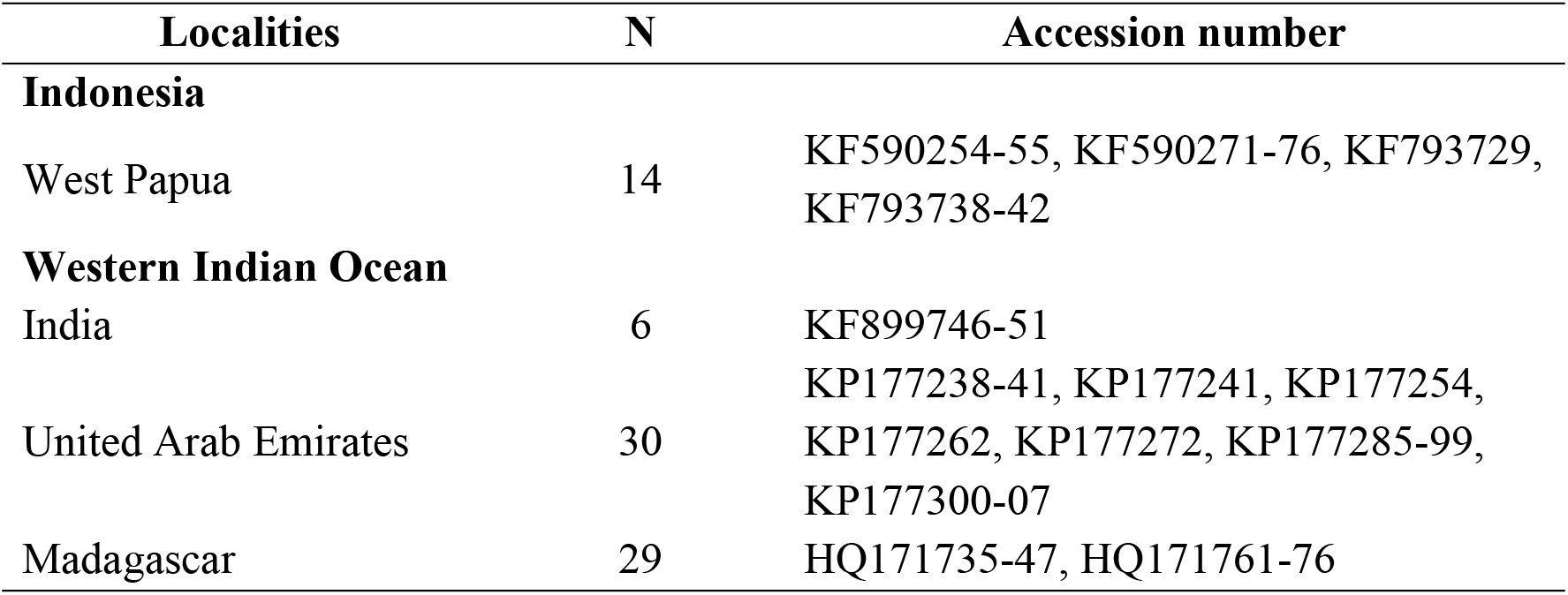
Localities, total number (n), and accession number of COI gene sequences of *S. lewini* in Western Papua (Indonesia), India, United Arab Emirates and Madagascar (Western Indian Ocean) retrieved from GenBank

#### Population structure

An analysis of molecular variance (AMOVA) and Wright’s fixation index (*F*_ST_) [26] were conducted for three major group: 1) within and among four population in Indonesia, 2) within and among three population in Western Indian Ocean, and 3) comparation between population from Indonesia and Western Indian Ocean using the Arlequin v.3.5 program (setting up 1000 permutations, α= 0.05 for significance level threshold) to estimate overall extent of genetic variation and differentiation level in Indonesia and Western Indian Ocean. Population differentiation and its significance between sampling sites were also calculated with pairwise estimates. The value of *F*_ST_ is a range of zero to one, low level of differentiation was implied from the closer of *F*_ST_ value to zero, and otherwise the high level will be closer to one [27–28].

#### Genetic connectivity

A haplotype network was constructed with a median-joining method in the Network v5.1.1.0 program [29] for all haplotypes found in Indonesia and Western Indian Ocean to obtain haplotype connectivity in broader spatial connection in the regional area in the Indian Ocean. Distribution of haplotypes for each location was also provided in a proper map to show clear distribution of haplotypes and genetic connectivity among populations.

## Results

### Genetic Diversity

A total of 179 sequences of the 594 bp mitochondria CO1 gene of *S. lewini* obtained from three sampling sites (Aceh, Balikpapan, Lombok), and additional sequences of West Papua and West Indian Ocean region were generated a total of 11 haplotypes variation with 40 polymorphic loci (Table 2). All the sequences of *S. lewini* obtained from this study were deposited in BOLD System with Barcode Index Number (BIN) registry of BOLD:AAA2403.

**Table 2.**
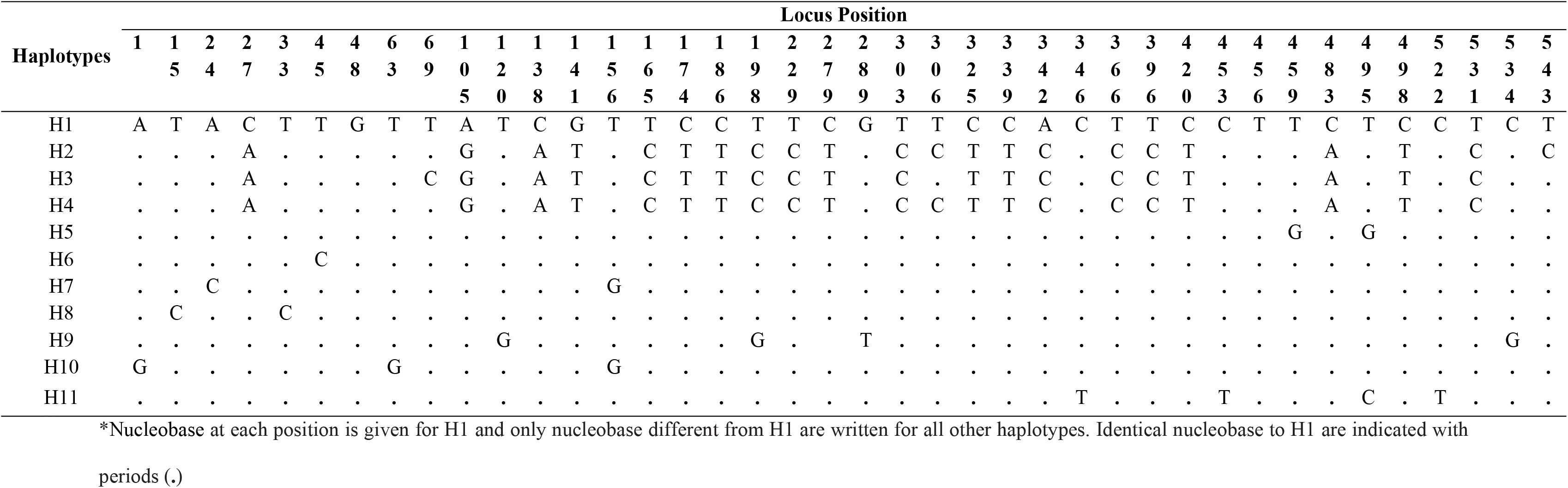
Forty polymorphic loci of 11 haplotypes from 179 sequences of the mitochondrial COI gene of *S. lewini* from four localities in Indonesia and three localities in Western Indian Ocean

Comparison of genetic diversity of *S. lewini* based on haplotype and nucleotide diversity was varied in value (Table 3). The haplotype diversity (Hd) among localities in Indonesia was ranged from 0.1429-0.6683 and nucleotide diversity (π) was ranged from 0.005-0.0198. The highest value of genetic diversity in Indonesia was showed by population in Aceh (Hd= 0.6683; π= 0.0198), followed by Balikpapan population which lower in haplotype and nucleotide diversity (Hd= 0.6460; π= 0.0020), whereas the lowest was West Papua (Hd= 0.1429, π= 0.0005). Population of *S. lewini* in Lombok also had a fairly low genetic diversity value (Hd= 0.3621; π= 0.0010). However, overall genetic diversity in Indonesia relatively high in value (Hd= 0.7171; π= 0.0126). On the other hand, the value of genetic diversity of Western Indian Ocean was low in average with ranged of 0.000-0.4667. The overall diversity in Western Ocean also low in value (Hd= 0.2322; π= 0.0010).

**Table 3.** Genetic diversity of *S. lewini* based on sample size (n), haplotype number (Hn), haplotype diversity (Hd), and nucleotide diversity (π) from each population sites in Indonesia and Western Indian Ocean region

| Population | n | Genetic Diversity |  |  |
| --- | --- | --- | --- | --- |
|  |  | Hn | Hd | Π |
| Indonesia |  |  |  |  |
| Aceh (ACH) | 41 | 4 | 0.6683 | 0.0198 |
| Balikpapan (BPN) | 30 | 3 | 0.6460 | 0.0020 |
| Lombok (LOM) | 29 | 3 | 0.3621 | 0.0010 |
| West Papua (WEP) | 14 | 2 | 0.1429 | 0.0005 |
| Overall Indonesia | 114 | 4 | 0.7171 | 0.0126 |
| Western Indian Ocean |  |  |  |  |
| India (IND) | 6 | 1 | 0.0000 | 0.0000 |
| United Arab Emirates (UAE) | 30 | 8 | 0.4667 | 0.0022 |
| Madagascar (MDG) | 29 | 1 | 0.0000 | 0.0000 |
| Overall Western Indian Ocean | 65 | 8 | 0.2322 | 0.0010 |

### Population structure

The analysis of fixation index (*F*_ST_) and significant P-value between and within four *S. lewini* populations (ACH, BPN, LOM, and WEP) in Indonesia and three population (IND, UAE, and MDG) were showed in Table 4. The overall value of *F*_ST_ in Indonesia was high and significant (*F*_ST_= 0.442; P-value= 0.000) which indicated that there were multiple subdivition in Indonesia. Low *F*_ST_ value and no significant different was exhibited by population from Western Indian Ocean (*F*_ST_= −0.0131; P-value= 0.6011), while comparative population structure between Indonesia and Western Indian Ocean showed significant differentiation (*F*_ST_= 0.7403; P-value= 0.0000). The pairwise *F*_ST_ between population in four location in Indonesia and comparison with Western Indian Ocean population was showed in Table 5. The overall pairwise analysis by distance method indicated significant differentiation among four population in Indonesia, except between BPN and LOM which showed fairly low *F*_ST_ value and no significant P-value. Furthermore, among population in Indonesia ACH population was tent to closer to WIO population with lower *F***_ST_** value rather than *F***_ST_** range value of other localities in Indonesia to Western Indian Ocean.

**Table 4.** Analysis of molecular variance (AMOVA) for percentage of variation (%), *F*_ST_ value and significant level (P-value) of *S. lewini* for Indonesian population, Western Indian Ocean population and between Indonesian and Western Indian Ocean population

| Source of variation | df | Percentage of variation (%) | $F_{ST}$ value | P-value |
| --- | --- | --- | --- | --- |
| Indonesia |  |  |  |  |
| Among Population | 3 | 44.15 | 0.4415 | 0.000 |
| Within Population | 110 | 55.85 |  |  |
| Total | 113 |  |  |  |
| Western Indian Ocean |  |  |  |  |
| Among Population | 2 | -1.31 | -0.0131 | 0.6011 |
| Within Population | 62 | 101.31 |  |  |
| Total | 64 |  |  |  |
| Indonesia vs Western Indian Ocean |  |  |  |  |
| Among Population | 1 | 74.04 | 0.7403 | 0.0000 |
| Within Population | 177 | 25.96 |  |  |
| Total | 178 |  |  |  |

**Table 5.** Pairwise *F*_ST_ value (below diagonal) and P-value (above diagonal) between *S. lewini* population in Aceh (ACH), Balikpapan (BPN), Lombok (LOM), West Papua (WEP), and Western Indian Ocean (WIO)

| Sample Sites | ACH | BPN | LOM | WEP | WIO |
| --- | --- | --- | --- | --- | --- |
| **ACH** | - | 0.0000 | 0.0000 | 0.0000 | 0.0000 |
| **BPN** | 0.4271 | - | 0.0898 | 0.0010 | 0.0000 |
| **LOM** | 0.4377 | 0.0435 | - | 0.0000 | 0.0000 |
| **WEP** | 0.3981 | 0.4945 | 0.7355 | - | 0.0000 |
| **WIO** | 0.5094 | 0.9647 | 0.9731 | 0.9750 | - |

### Genetic Connectivity

There were two main group of haplotypes that show by haplotype network analysis (Fig 1), hereafter referred to as Clade A and Clade B. Clade A consisted of H1 found in multiple region in Aceh, India, United Arab Emirates, and Madagascar, while H5, H6, H7, H8, H9, H10, H11 were originated only from United Arab Emirates. Clade B consisted of three haplotypes (H2, H3, H4) which spread evenly in Indonesia, though there were predominant haplotype found in every single region. Aceh, Balikpapan, Lombok, and West Papua was predominated by H1, H2, H4, and H3 respectively (Fig 2).

**Fig 1.**
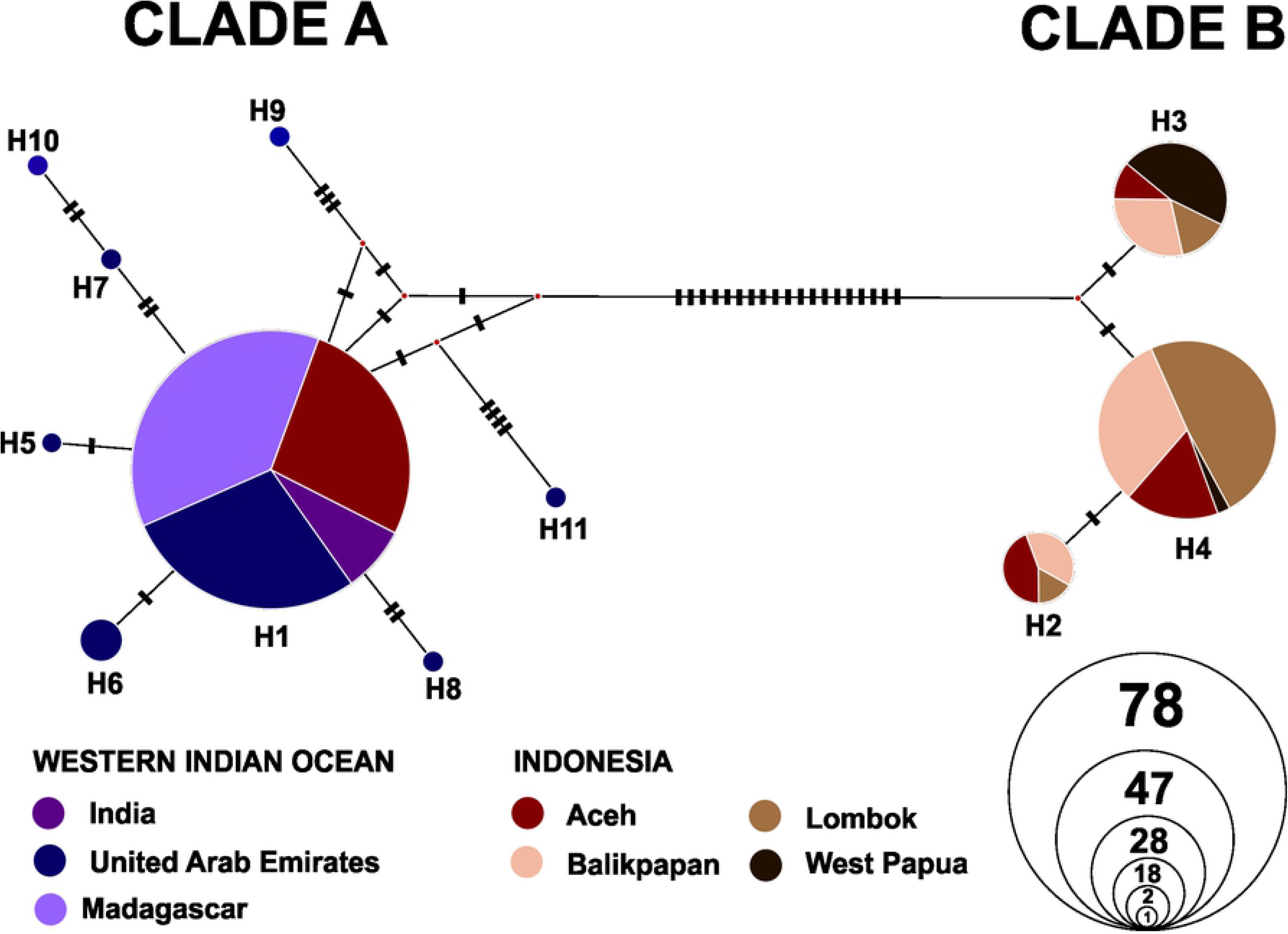
Haplotype network of *S. lewini* (n= 179) population in Indonesia and Western Indian Ocean region, constructed with Median Joining method.

**Fig 2.**
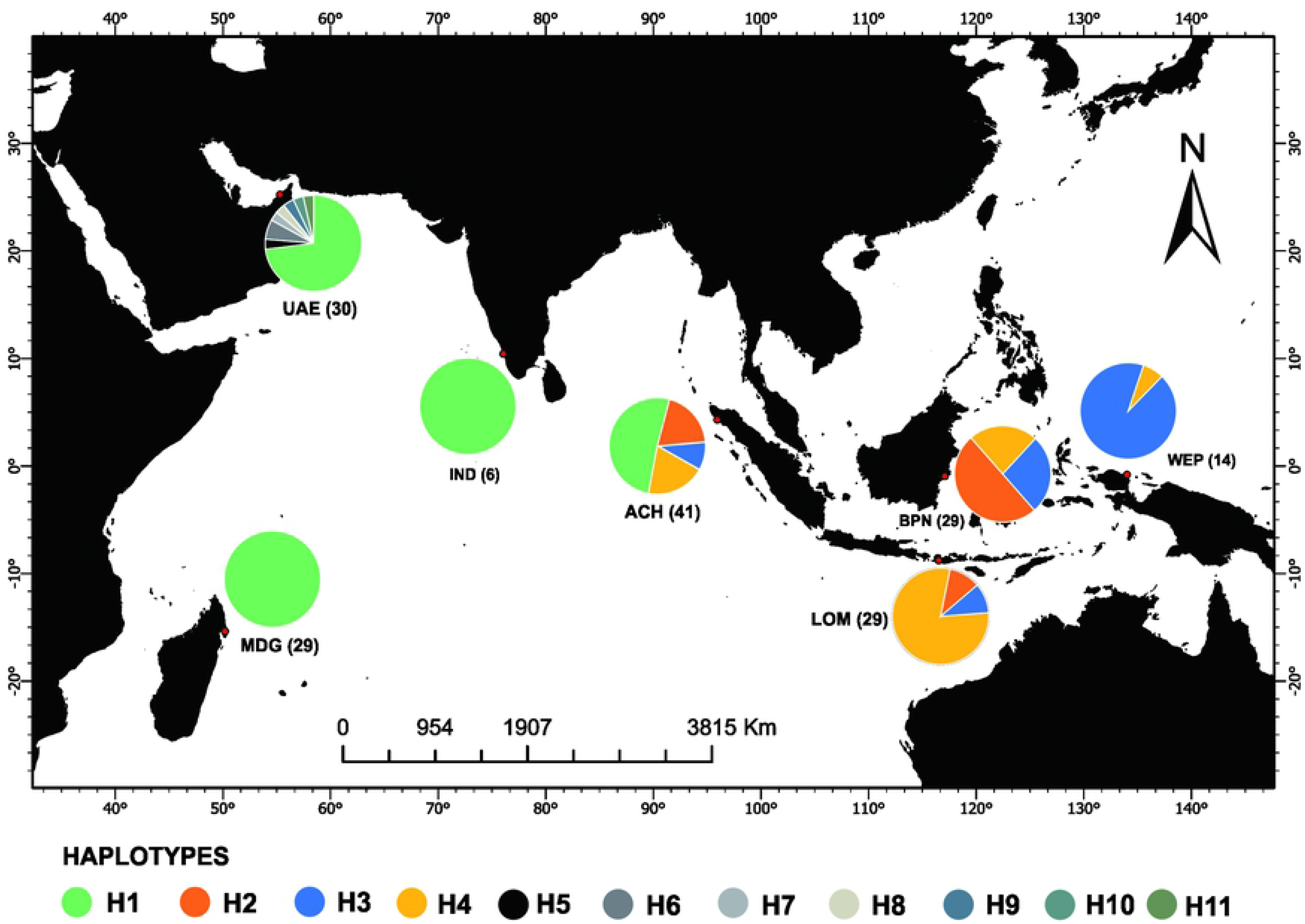
Distribution of 11 haplotypes of *S. lewini* population from Indonesia and Western Indian Ocean in regional scale.

## Discussion

### Genetic diversity

The overall genetic diversity both at the haplotype and nucleotide level of *S. lewini* was relatively high for population in Indonesia. This finding is consistent with the result of Ovenden *et al.* [14] with mitochondria control region of *S. lewini* from three localities in Indonesia including Lombok and high genetic diversity of *S. lewini* in global scale [13]. The scalloped hammerhead sharks are high migratory species with a wide distribution in tropical and warm-temperate waters and possibly move across oceanic waters up to 1500 km far [9]. In principle, with migration capability and broad ecological niches, this species tends to have higher genetic diversity than narrow ecological niches species [30].

High genetic diversity values are commonly related to large population size [31], also promotes by some factors such local population sizes, fast generations times [32], high nucleotide substitution rate [33], and the high gene flow between geographical distant populations.

The finding of relatively high genetic diversity of *S. lewini* in Indonesia appear inconsistent with general assumption that overexploitation as both target fishing and bycatch have declined the scalloped hammerhead shark worldwide population [34], however, this could be relevant with the fact occurred in Lombok. Lower genetic diversity evidence occurred in Lombok probably driven by continuous fishing pressure. *S. lewini* were the top three targeted sharks landed in Tanjung Luar fishing port, Lombok and have faced high fishing pressure more than 40 years that lead to overfishing with exploitation rate (E) reached 0.59 recently [35–36].

Furthermore, according to fisheries global information system statistic of FAO [37] the hammerhead sharks (Sphyrnidae) including *S. lewini* are one of the important sharks which highly exploited in the last two decades which estimated rapidly increase in global capture since 220 ton per year in 1985 reaching approximately 10,362 tons per year in 2016. In the same period the capture level in Indonesia also significantly increase reaching approximately 1,492 tons in 2016 with the fact that Indonesia was in the highest volume of shark and rays caught on the global catches reported in 2000-2011 [38].

Clarke *et al*. [39] was reported similar finding regarding global mitochondrial DNA of silky shark, *Carcharhinus falciformis* with high genetic diversity under circumstance of overexploitation. Elasmobranch possible to have adaptability with environmental and anthropogenic stress which causes genetic bottlenecks because of their particular life histories [40]. Moreover, population decline caused by fishery activities recently might be insufficient in time to reduce genetic diversity particularly for long life species (13–20 years) including S. lewini [41–42].

However, indeed declining large population continuously, as was expected occur with *S. lewini* population in Lombok, had positive correlation with loss of genetic diversity and driven a population bottleneck [42–43] as revealed by Pinsky and Palumbi [44] in some marine fish meta-analysis.

### Population structure

Our finding indicated homogeneity among *S. lewini* population from Balikpapan and Lombok. Pairwise *F*_ST_ analysis detected no significant genetic differentiation in these two contiguous spatial distant. This finding complements with previous studies conducted in Indo-Australia waters. Ovenden *et al.* [14] reported similar evidence for mitochondrial control region on population structure with no differentiation for two population of *S. lewini* between Indonesian population including Lombok and northern Australian. This pattern of single stock population seems to suggest that these localities are migration zone of *S. lewini* and reproductive movement might occur in coastal area of these locations though there are strong Indonesian throughflow current between Kalimantan and Sulawesi island. Adults *S. lewini* are highly migratory with large body size that supports of high dispersal capacity of this species in hence possibly to control that geographical barrier. Otherwise, we observed different result with comparison among these two population in central Indonesia with population in Aceh (western Indonesia) and West Papua (eastern Indonesia) which exhibited strong genetic differentiation. These regions were spatially separated by long distance with the possibility of more complex barriers both inter island barrier and anthropogenic factors of high commercial and artisanal fishing pressure such along southern and northern Java.

### Genetic connectivity

We found an interesting finding regarding haplotype sharing between population in Indonesia and Indian Ocean population. H1 was the unique haplotype found in Aceh, Indonesia and this haplotype was correlated with the most common haplotype in three locations of western Indian Ocean region from India, United Arab Emirates, and Madagascar which could indicated that there was connectivity between population from Aceh and western Indian Ocean. Schott and McCreary [45] described Indian Ocean seasonal current carried water from north Sumatera through Arabian Sea to Madagascar which was possible as connectivity channel of two regions with thus allowing *S. lewini* to migrate and lead to cross breeding. Similar predominant haplotype (H1) of *S. lewini* from Aceh with population from India, United Arab Emirates, and Madagascar were evidence of genetic sharing process between population in Indian Ocean region. However, the *S. lewini* population from Balikpapan, Lombok and West Papua was indicated isolation from population in western Indian Ocean and shared haplotype network only in eastern Indonesia water.

### Conservation implication

The overall high value of genetic diversity of *S. lewini* population in Indonesia suggests this species has not experienced a genetic loss and bottleneck to face of exploitation pressure. However, lower genetic diversity of *S. lewini* in Lombok and West Papua indicated a higher risk of genetic diversity loss that probably as the result of high fisheries pressure. Our genetic assessment of *S. lewini* from four localities which revealed a single stock exist between Lombok and Balikpapan, and separated stock for Aceh and West Papua indicating that management of this species should occur on stock-based approach at least on three mitochondrial-stock conservation management unit. A complex and limited connectivity population of *S. lewini* that existed in population in Indonesia, and between Western Indian Ocean indicating the importance to promote collaborative management strategic between Indonesian management agencies, and also with West Indian Ocean agencies in regional scales.

## Conclusion

This study has shown important findings regarding population structure of *S. lewini* in Indonesia with high genetic diversity and three significant subdivision that indicated capability of population to adapt to rapid environmental change and pressure including fishing activities. However, lower genetic value in Lombok and West Papua should still be considered. Genetic sharing which detected between population in Indonesia and Western Indian Ocean proved connectivity between population in Indonesia and also connecting with Western Indian Ocean population. Collaborative action across region is needed to promote sustainable management and conservation purpose both in Indonesia and Western Indian Ocean in regional scale.

## Acknowledgement

This research was supported by Wildlife Conservation Society (WCS)-Indonesia, in collaboration with Faculty of Fisheries and Marine Sciences IPB University. Sutanto Hadi received a scholarship from Indonesia Endowment Fund for Education (LPDP). We also appreciated to BPSPL Satker Balikpapan, Head of PPI Manggar Balikpapan, and also Marine Biodiversity and Biosystematic Lab (BIODIVSI) fellows for their help during research, and to reviewers who have provided highly constructive suggestions for better writing of the paper.

